## Supplemental Figures and Tables for "Genome-scale analysis of interactions between genetic perturbations and natural variation"

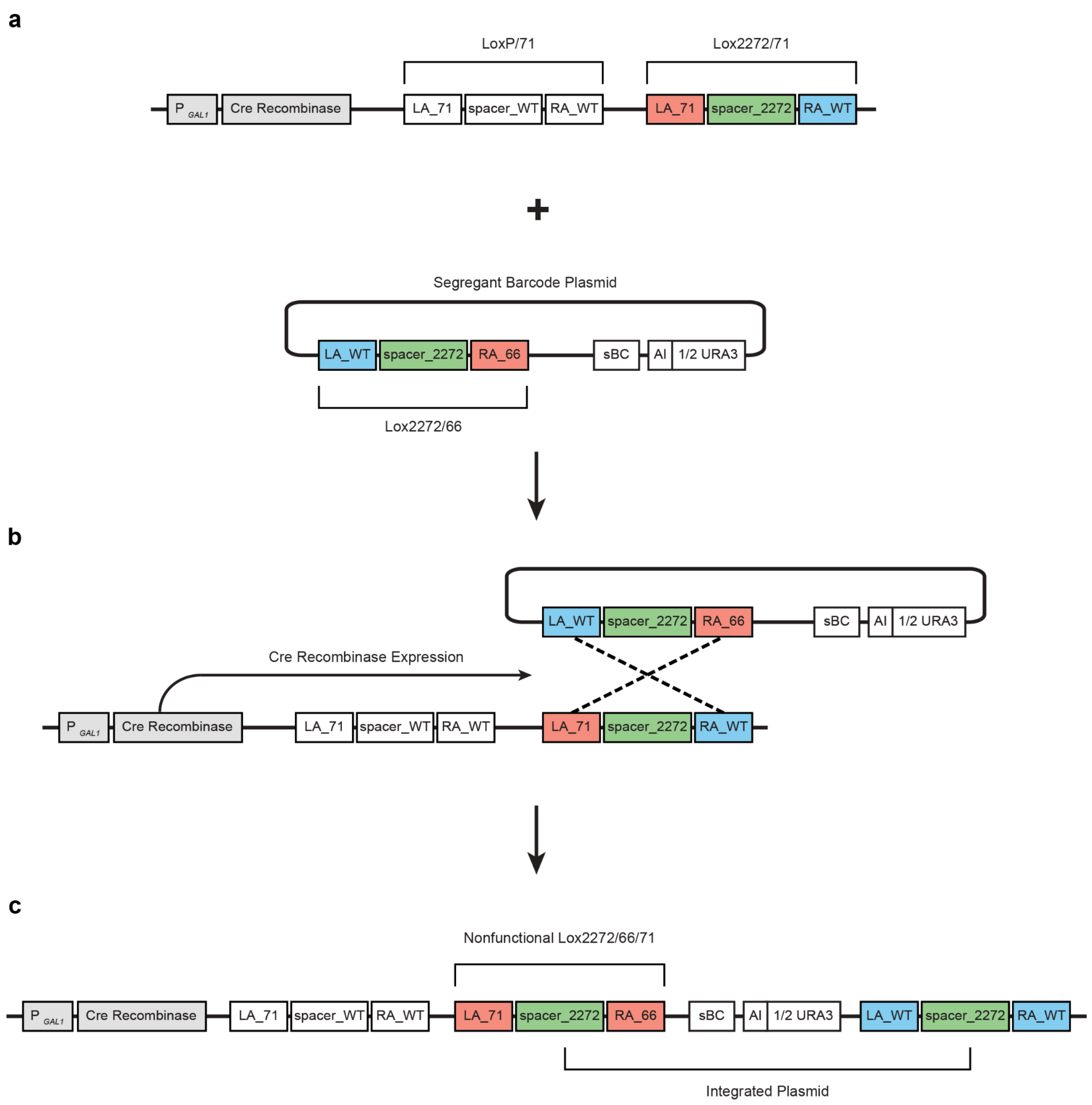


**Supplemental Figure 1: Landing Pad design.** a) Genomic landing pad before integration of any plasmids. LA_71 indicates a partial mutation on the left arm of the Lox site, which has no effect on its own. RA_66 indicates a similar partial mutation on the right arm. Lox2272/66 should only recombine with Lox2272/71. b) Expression of Cre recombinase causes the plasmid to be integrated into the genome in a predictable configuration. c) Recombination at the Lox2272 sites results in the plasmid contents being integrated into the chromosome. This also places both mutant arms on the same Lox site, which renders it nonfunctional and limits spontaneous popout of the plasmid.

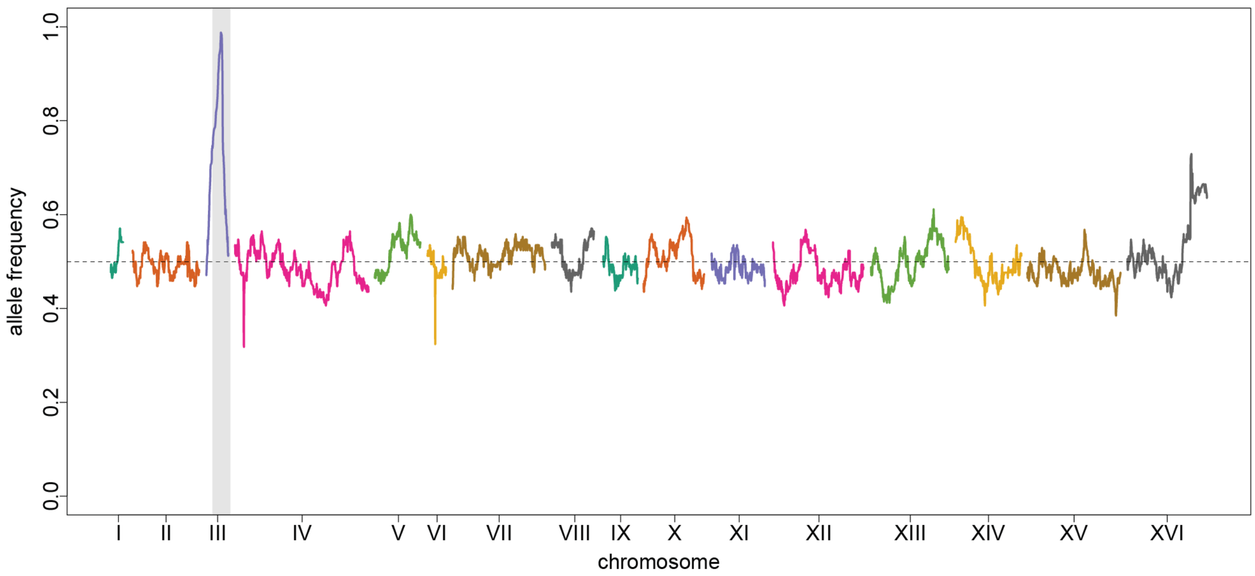


**Supplemental Figure 2: Allele frequencies across all genotypes.** The proportion of segregants carrying the 3S allele of a site is shown. The fixed region on Chromosome III is the mating locus, at which all segregants possess the 3S allele of MATalpha.

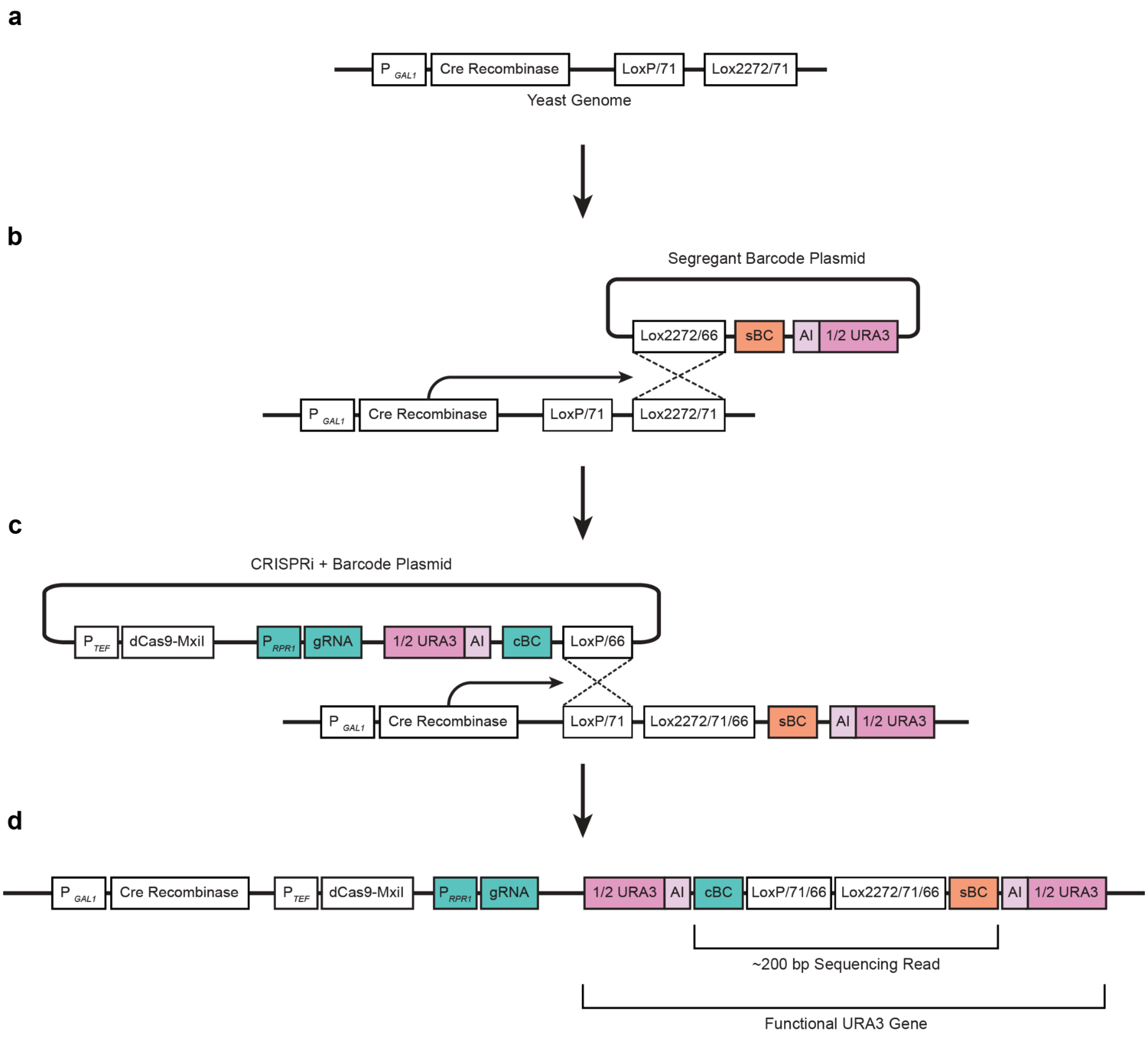


**Supplemental Figure 3: Plasmid design and double-barcoding scheme.** a) Genomic landing pad before integration of any plasmids. b) Integration of segregant barcode plasmid using the Lox2272 sites. A detailed visualization of this step is available in Supplemental Figure 1. c) Integration of barcoded CRISPRi plasmid library using the LoxP sites. d) Landing pad structure after CRISPRi plasmid integration. The *URA3* gene is only functional after CRISPRi plasmid integration, allowing for selection in SC - URA liquid media.

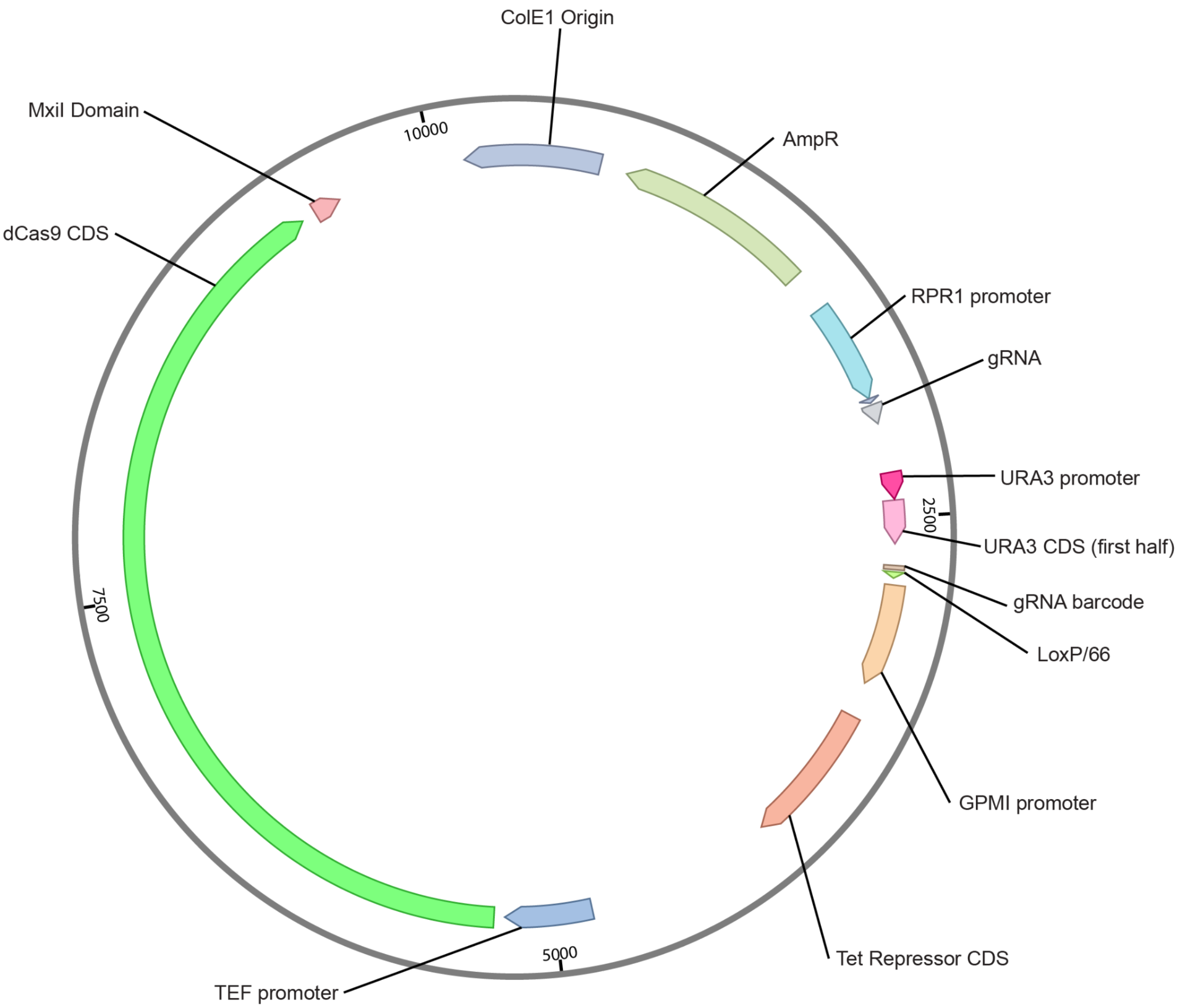


**Supplemental Figure 4: CRISPRi plasmid map.**

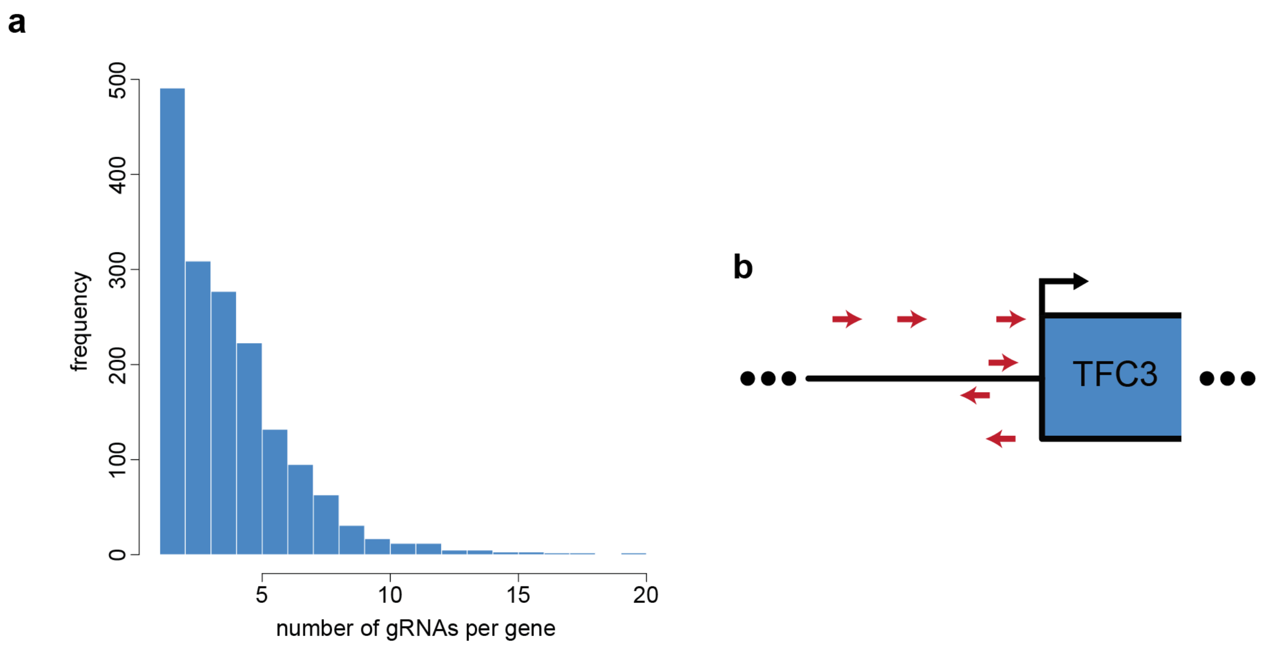


**Supplemental Figure 5: Multiple replicate gRNAs target the same gene.** a) Number of replicate gRNAs per gene (mean = 4.14, median = 4, sd = 2.65). b) An example of replicate gRNA positioning relative to a target gene. The blue box indicates the coding sequence of *TFC3/YAL001C*, with all gRNAs positioned around the promoter. Arrows indicate guide strand (plus/minus).

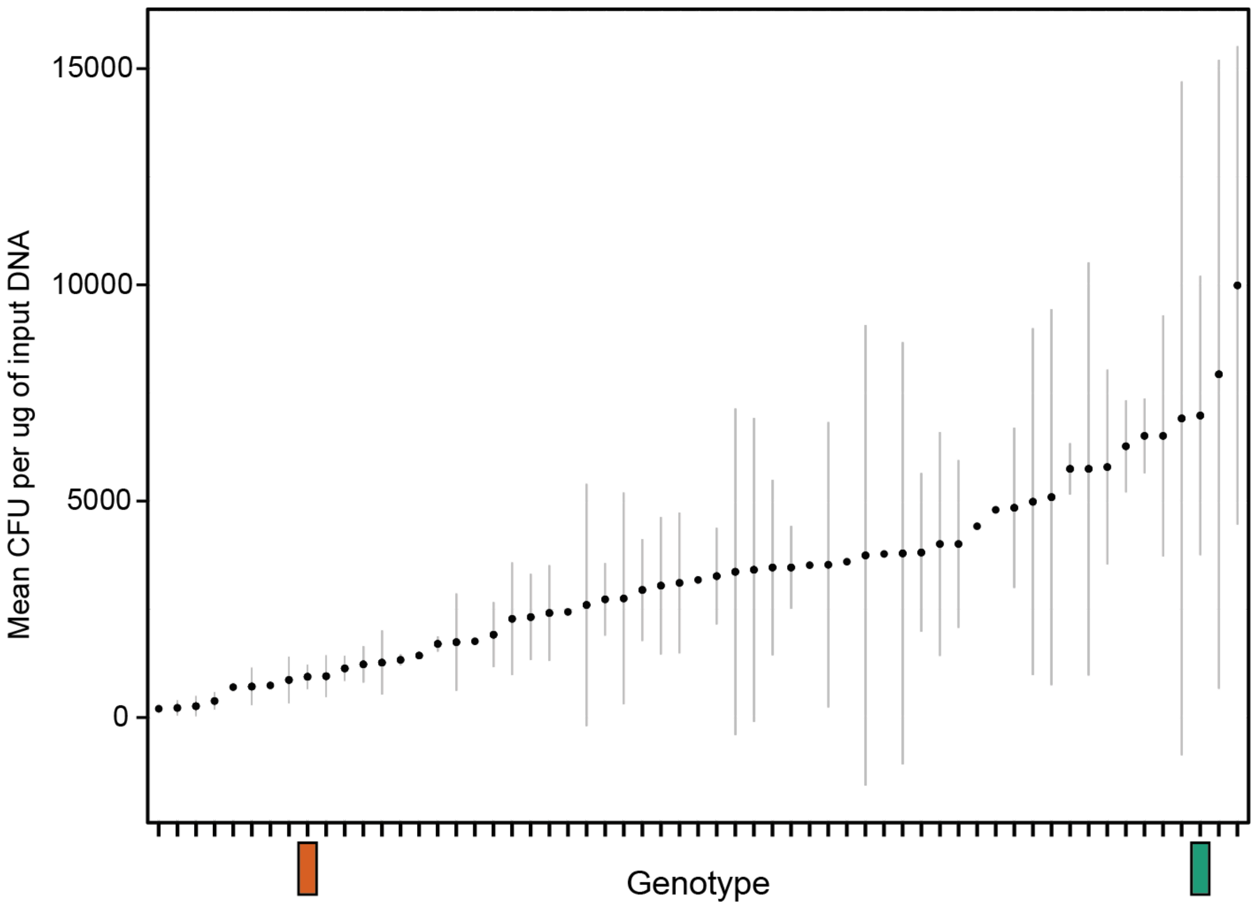


**Supplemental Figure 6: Transformation efficiency across 58 random segregants.** Segregants without error bars were not replicated. Dots represent means and error bars represent one standard deviation. The green bar indicates the BY parent, and the orange bar indicates the 3S parent.

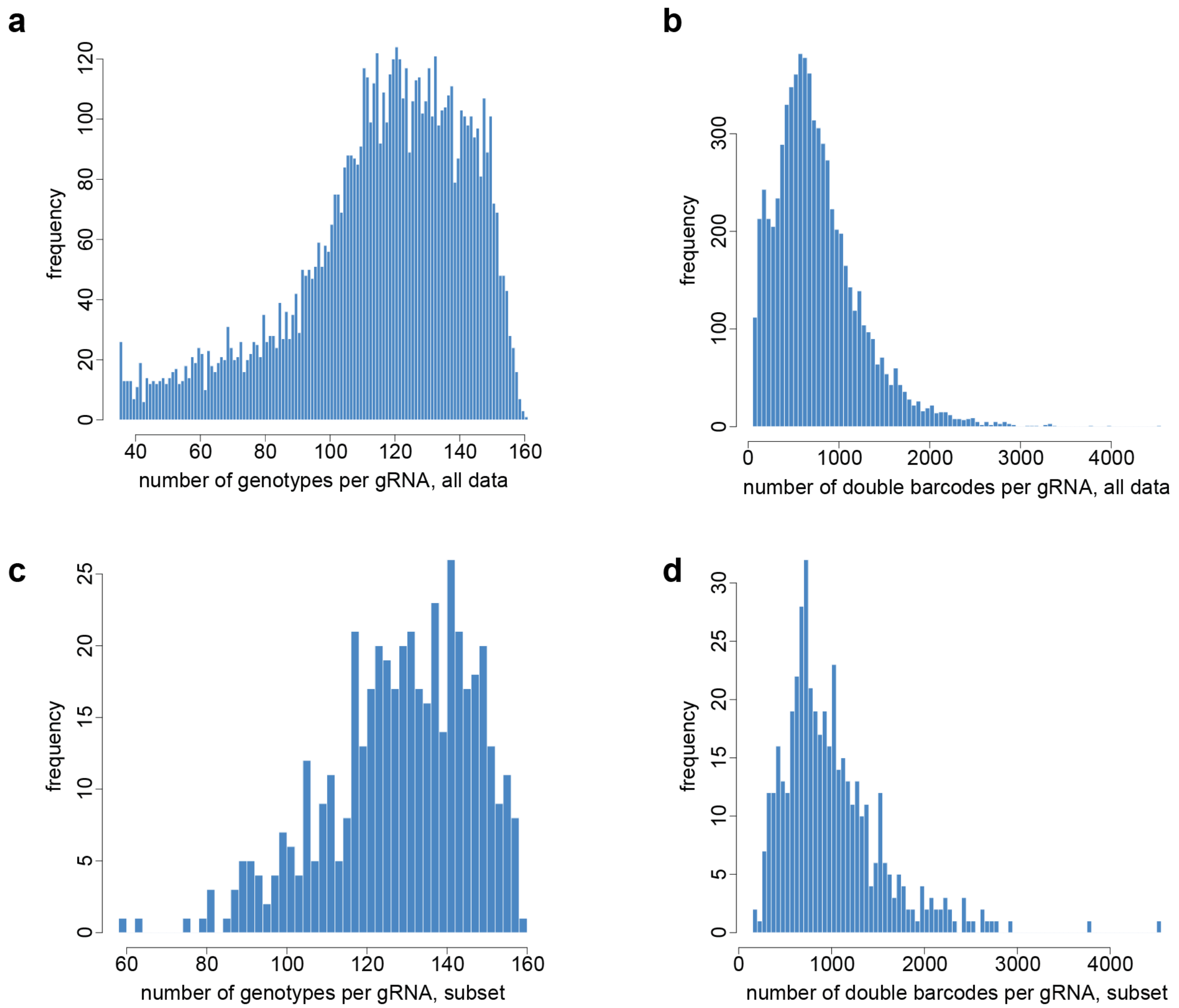


**Supplemental Figure 7: Distribution of data per gRNA.** a) Histogram of the number of genotypes per gRNA for all ~7,000 analyzed gRNAs (mean = 115.98, median = 120, sd = 26.84). b) Number of double barcodes per gRNA for all analyzed gRNAs (mean = 762.61, median = 674, sd = 474.65). c) Number of genotypes per gRNA for the 460 efficacious gRNAs used in downstream analyses (mean = 128.85, median = 131, sd = 18.14). d) Number of double barcodes per gRNA for gRNAs used in downstream analyses (mean = 1008.15, median  = 892, sd = 549.26).

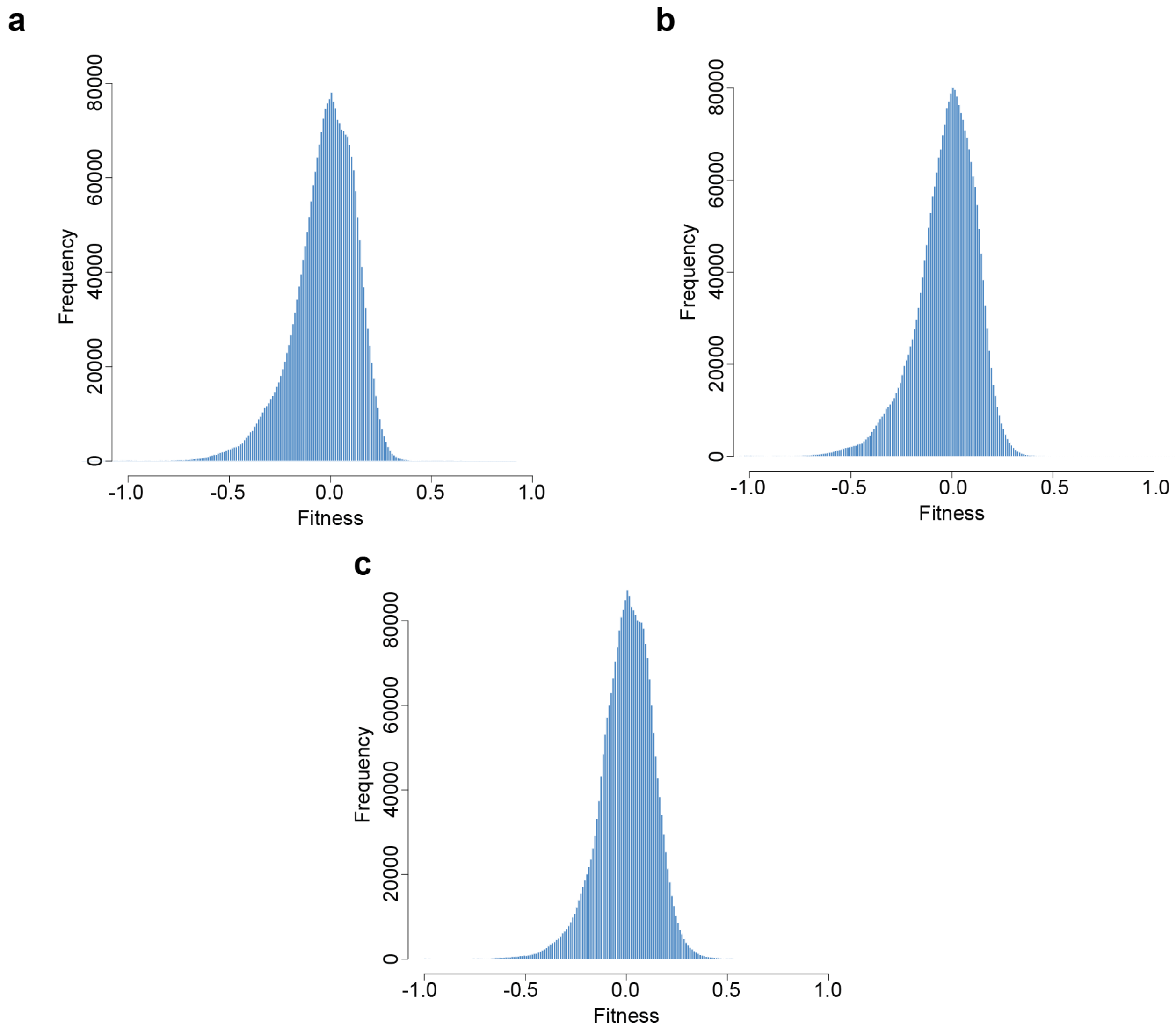


**Supplemental Figure 8: Distribution of fitness values from each assay.** a) Distribution of fitness values in ATC1. b) Distribution of fitness values in ATC2. c) Distribution of fitness values in CON.

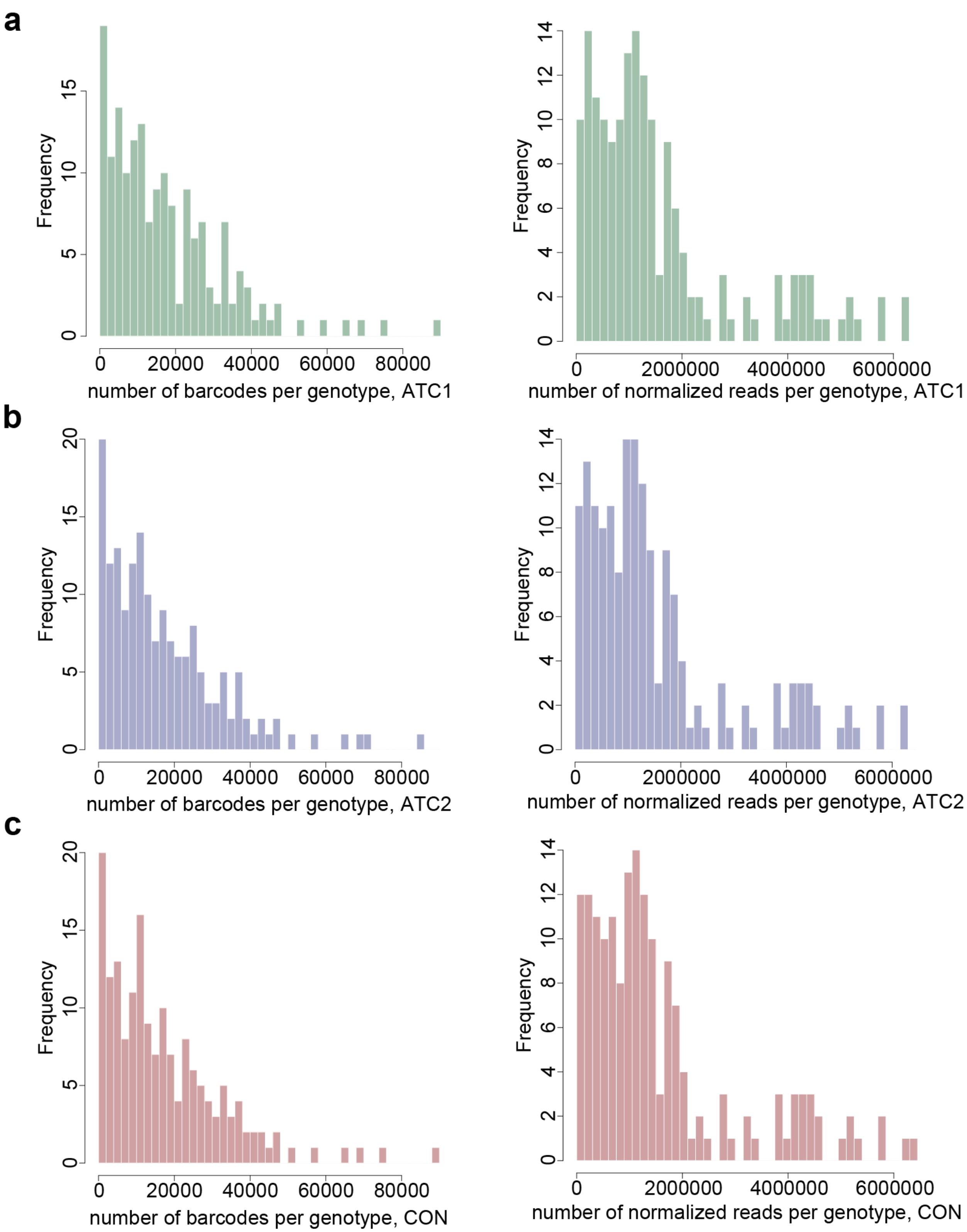


**Supplemental Figure 9: Distribution of barcodes and normalized reads per genotype in each flask.**

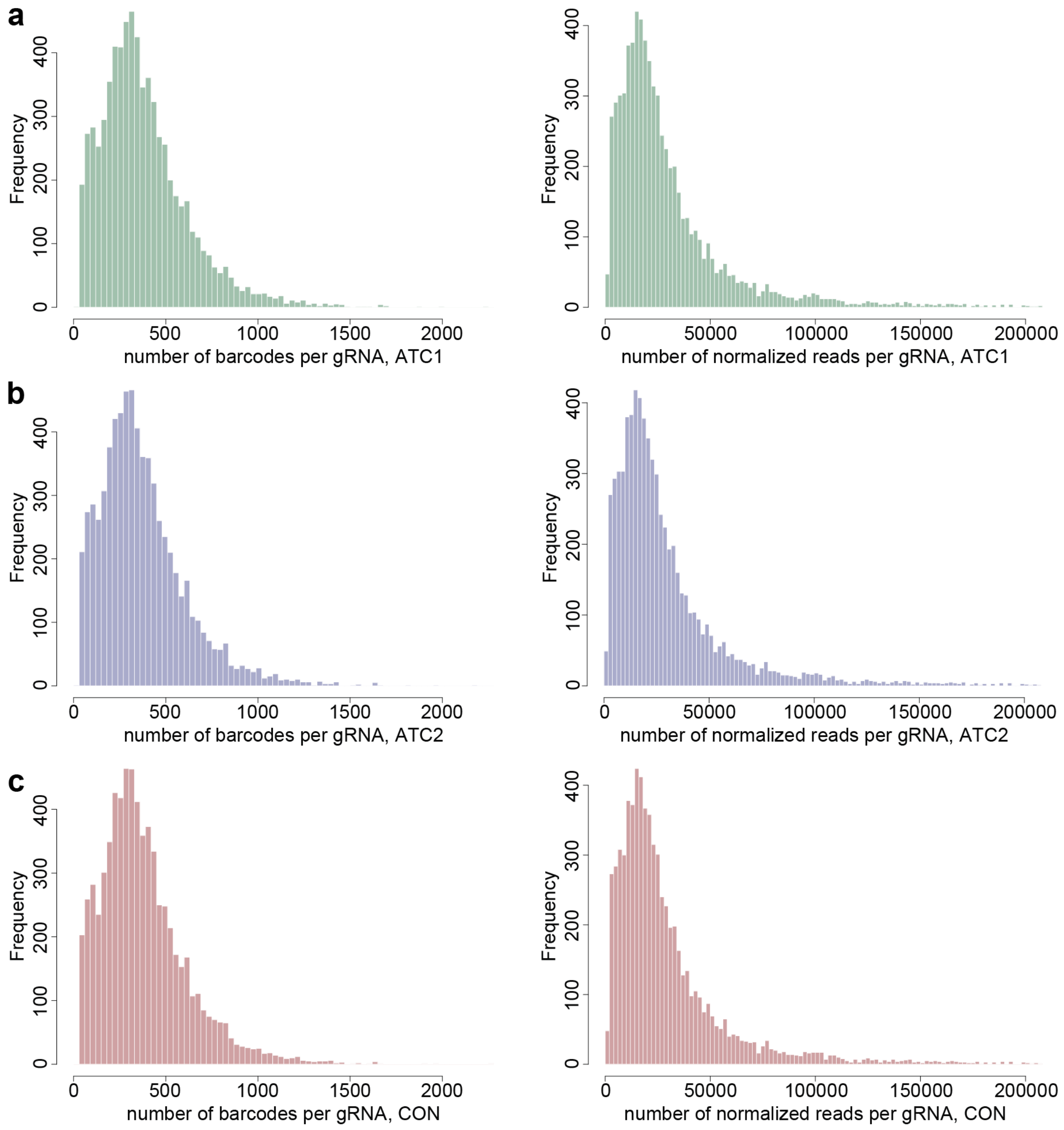


**Supplemental Figure 10: Distribution of barcodes and normalized reads per gRNA in each flask.** Normalized read histograms have extreme outliers removed for visibility (~1.5% of all gRNAs).

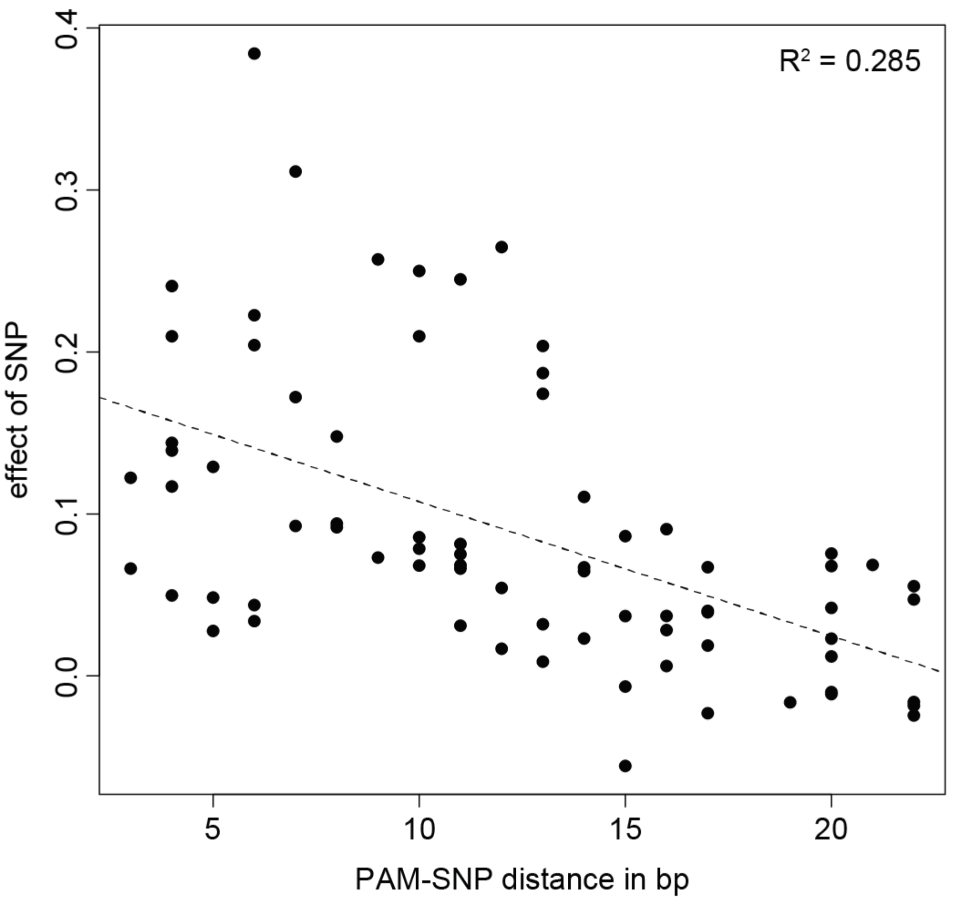


**Supplemental Figure 11: XY plot between the effect of SNPs that directly overlap gRNA binding sites and the distance between the gRNA's PAM site and that SNP.** The effect of the SNP is calculated as (effect of the guide in segregants with the BY allele) - (effect of the guide in segregants with the 3S allele). gRNAs perfectly match their targets in segregants with the BY allele. Simple linear regression $R^2 = 0.285$, pvalue = $9.9 \times 10^{-7}$.

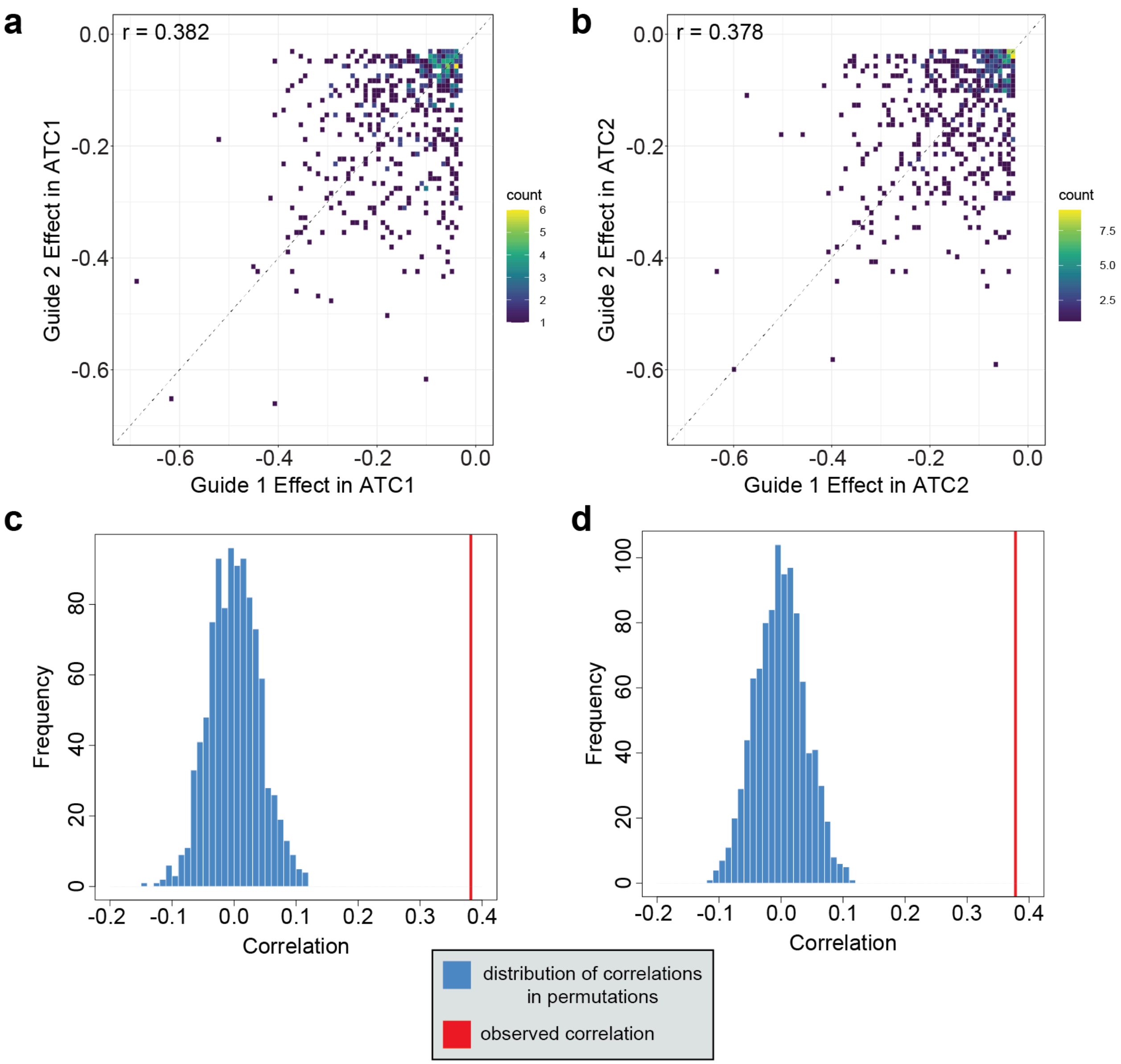


**Supplemental Figure 12: Effects of gRNAs that target the same gene.** gRNA effect was estimated by a mixed effects linear model (see Methods section "Identification and Quantification of Guide Effects"). If more than two gRNAs targeted the same gene, they were randomly assigned into pairs. The same subset of gRNAs was used for all following panels. a) X-Y plot showing the correlation between effects among gRNA pairs in the ATC1 condition (Pearson correlation = 0.382, pval = $5.61 \times 10^{-19}$). b) X-Y plot showing the correlation between effects among gRNA pairs in the ATC2 condition (Pearson correlation = 0.378, pval = $5.66 \times 10^{-20}$). c) Permutations for the correlation shown in (a). All gRNAs were randomly assigned into pairs, regardless of their targeted gene, and the same Pearson correlation was calculated. This process was repeated 1000 times, with the resulting distribution of correlations shown in blue. The value from the original data set is shown by the red line. d) Permutations for the correlation shown in (b).

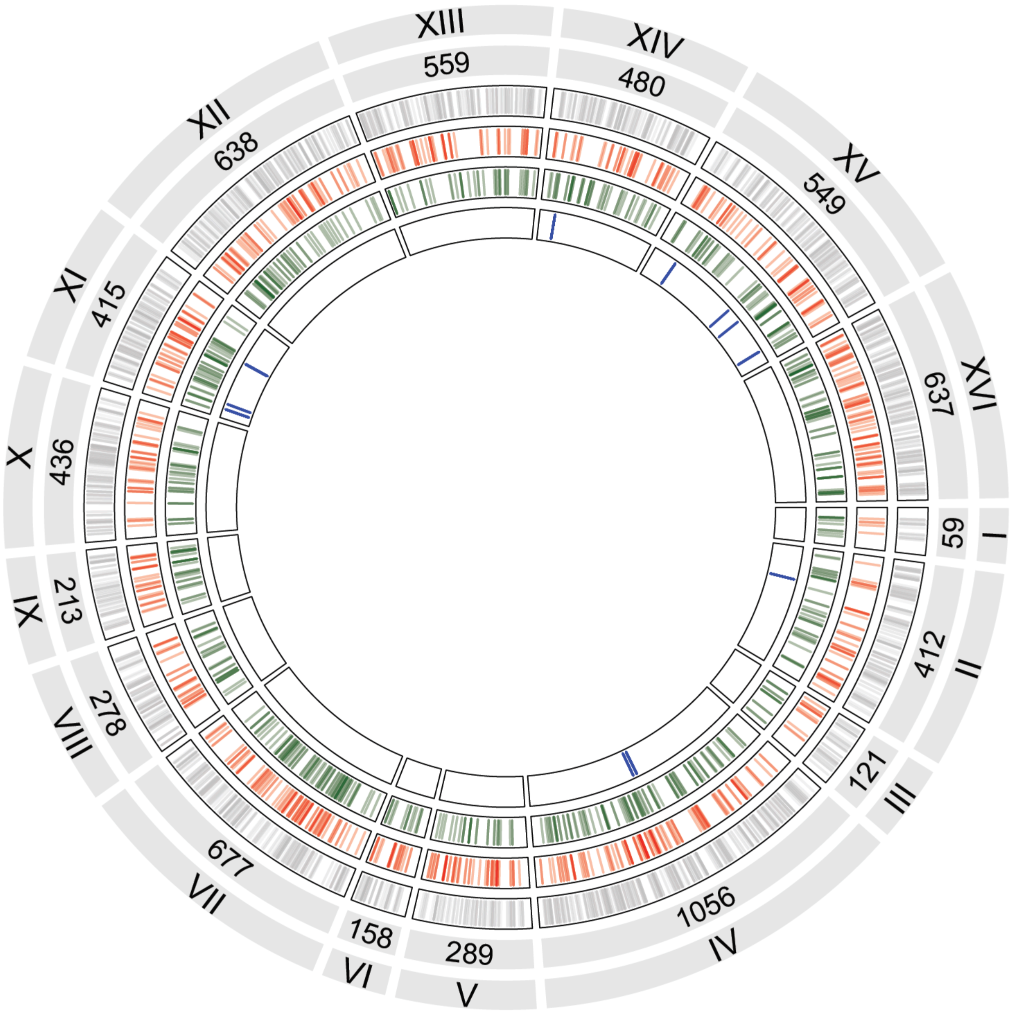


**Supplemental Figure 13: Location of all gRNAs across the genome.** Numbers show how many gRNAs in total are present on each chromosome. Gray lines indicate locations of gRNAs without mean effects. Red lines indicate locations of gRNAs with mean effects only. Green lines indicate gRNAs with background effects. Blue lines indicate control gRNAs.

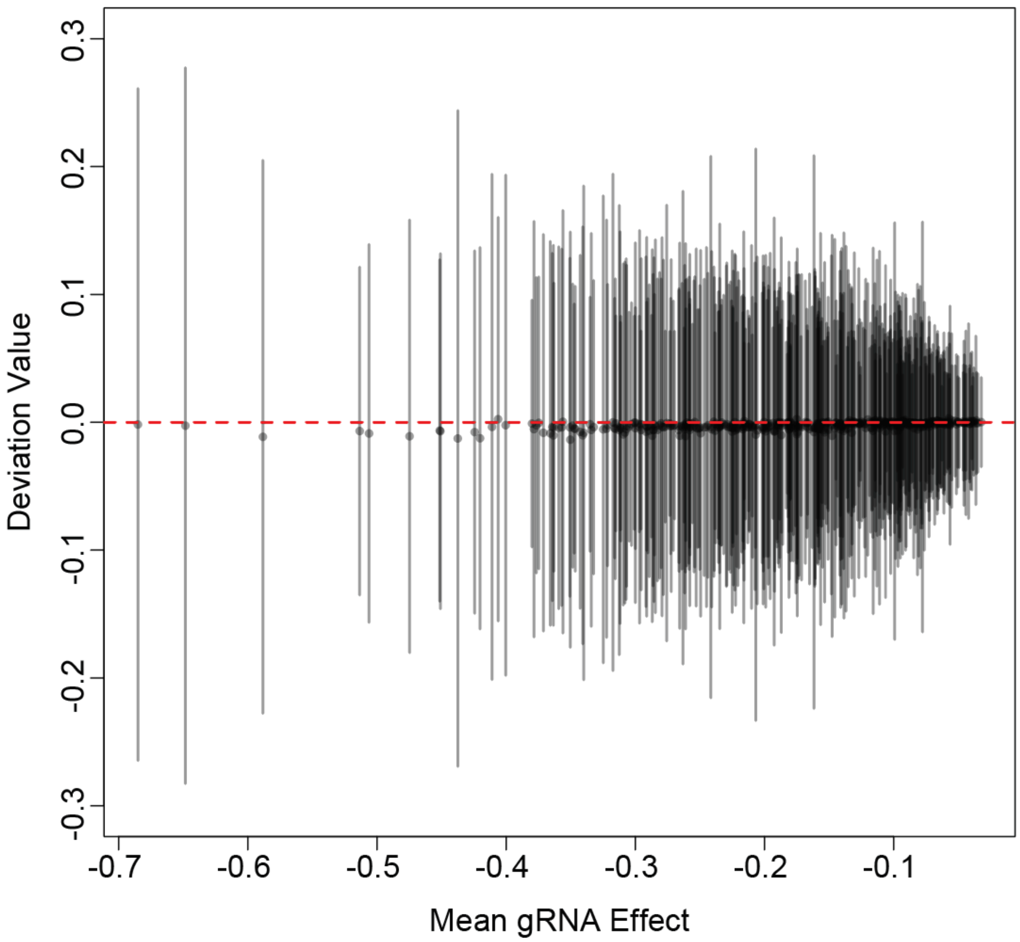


**Supplemental Figure 14: X-Y plot between gRNA effect and all deviation values for that gRNA.** Mean deviation value is shown by a point, with lines indicating two standard deviations. Each point represents a distinct gRNA-targeted gene with a background effect.

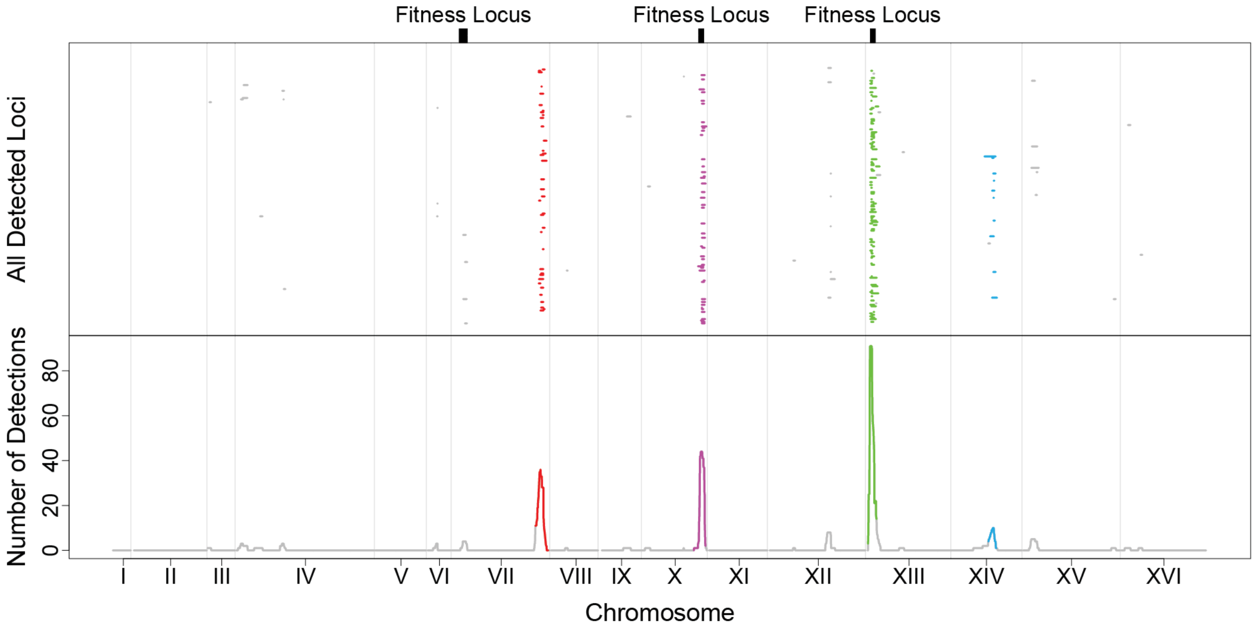


**Supplemental Figure 15: Linkage mapping results on deviation values for the replicate experimental flask ATC2.** Upper panel: Visualization of all 2x-$\log_{10}$(pval) drops for the gRNAs with significant peaks. Each row is a unique gRNA. Locations of major fitness loci are indicated by black bars. Lower panel: Number of overlapping 2x-$\log_{10}$(pval) drops at each nucleotide position along the genome. Regions where more intervals overlap than expected by chance (hubs) are shown in unique colors. Vertical lines indicate chromosome boundaries.

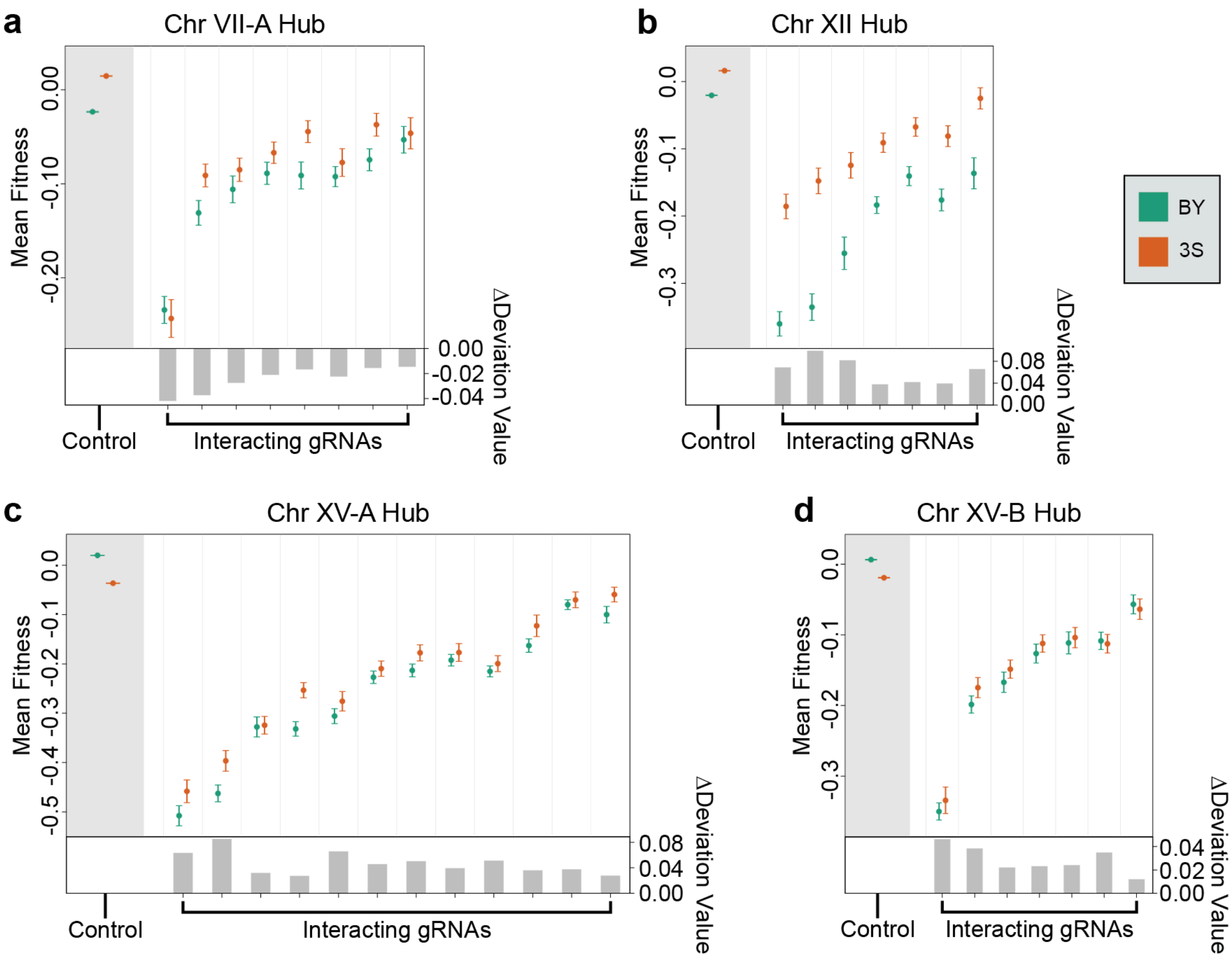


**Supplemental Figure 16: Potential hubs below the significance threshold used during analysis.** Hubs were categorized here if they passed the significance threshold of 9 overlapping confidence intervals in only one of the two conditions (ATC1 or ATC2), or if at least 5 overlapping confidence intervals were present in at least one condition. a) Effect of a locus near the 5' end of chromosome VII, at position 119852 and 39054 bp in width. This locus is distinct from the main Chr VII hub, and was detected as a significant locus during the linkage mapping on baseline fitness. This locus was detected only in ATC1. b) Average effect of the locus on chromosome XII, at position 662627 and 27942 bp in width. c) Average effect of the locus near the 5' end of chromosome XV, at position 147540 and 59773 bp in width. This locus was above the significance threshold only in ATC1. d) Average effect of the second locus on chromosome XV, at position 1011045 and 11091 in width. This locus was only detected in ATC1.

**Supplemental Tables**

| Fitness Assay | Time Point | Total Raw Reads |
|---|---|---|
| ALL | 0 | 460428002 |
| ATC1 | 1 | 392520148 |
| ATC1 | 2 | 399527009 |
| ATC1 | 3 | 311268392 |
| ATC1 | 5 | 365532111 |
| ATC2 | 1 | 387400026 |
| ATC2 | 2 | 354986473 |
| ATC2 | 3 | 336666459 |
| ATC2 | 5 | 376377113 |
| CON | 1 | 333135885 |
| CON | 2 | 349830935 |
| CON | 3 | 302795126 |
| CON | 5 | 401794794 |

**Supplemental Table 1:** Total raw reads used for fitness estimation. Reads were counted just before performing normalization and running PyFitSeq (see Methods section "Fitness Estimation" for more detail).

| Time Point Set 1 | Time Point Set 2 | Pearson Correlation |
|---|---|---|
| T0-T1-T2-T3-T5 | T0-T1-T2 | 0.691 |
| T0-T1-T2-T3-T5 | T0-T1-T2-T3 | 0.824 |
| T0-T1-T2-T3-T5 | T0-T1-T2-T3-T5-T7 | 0.933 |
| T0-T1-T2-T3-T5 | T1-T2-T3-T5 | 0.888 |
| T0-T1-T2-T3-T5 | T2-T3-T5-T7 | 0.704 |
| T0-T1-T2-T3-T5 | T1-T2-T3-T5-T7 | 0.837 |
| T0-T1-T2 | T0-T1-T2-T3 | 0.822 |
| T0-T1-T2 | T0-T1-T2-T3-T5-T7 | 0.613 |
| T0-T1-T2 | T1-T2-T3-T5 | 0.509 |
| T0-T1-T2 | T2-T3-T5-T7 | 0.244 |
| T0-T1-T2 | T1-T2-T3-T5-T7 | 0.469 |
| T0-T1-T2-T3 | T0-T1-T2-T3-T5-T7 | 0.767 |
| T0-T1-T2-T3 | T1-T2-T3-T5 | 0.678 |
| T0-T1-T2-T3 | T2-T3-T5-T7 | 0.495 |
| T0-T1-T2-T3 | T1-T2-T3-T5-T7 | 0.651 |
| T0-T1-T2-T3-T5-T7 | T1-T2-T3-T5 | 0.833 |
| T0-T1-T2-T3-T5-T7 | T2-T3-T5-T7 | 0.788 |
| T0-T1-T2-T3-T5-T7 | T1-T2-T3-T5-T7 | 0.906 |
| T1-T2-T3-T5 | T2-T3-T5-T7 | 0.729 |
| T1-T2-T3-T5 | T1-T2-T3-T5-T7 | 0.936 |
| T2-T3-T5-T7 | T1-T2-T3-T5-T7 | 0.819 |

**Supplemental Table 2: Stability of fitness estimates across different subsets of time points.** In order to test how the choice of time points affects PyFitSeq fitness estimates, the software was provided with a variable number of input time points from the ATC1 fitness assay. For each unique set of time points, a fitness estimate was generated for every lineage. The Pearson correlation was obtained by comparing the fitness estimates for each lineage across two sets of input time points. The time point set "T0-T1-T2-T3-T5" was the one used in this study. All p-values were below $2.2 \times 10^{-16}$.

| gRNA Sequence | Chr | gRNA Start | gRNA End | PAM Position | SNP position | gRNA Effect |
|---|---|---|---|---|---|---|
| TGATGCATTCACTCTCAGTT | 12 | 805700 | 805719 | 805697 | 805710 | -0.0950996 |
| GAACTCTAATACCTTTAGTT | 2 | 691875 | 691894 | 691872 | 691880 | -0.0624023 |
| TTATCATAAACCAAGACCGG | 7 | 999243 | 999262 | 999240 | 999249 | -0.1315465 |
| AAAAGAGTAACCTAGGAAAT | 7 | 999219 | 999238 | 999216 | 999229 | -0.0916477 |
| AATATCACGATAGGACAAAT | 12 | 694205 | 694224 | 694202 | 694211 | -0.0356261 |
| AGGCCTGAAAAATTTCAAAG | 11 | 122400 | 122419 | 122422 | 122402 | -0.0863247 |
| GGAGAAAACTAAACGGGAAT | 7 | 894494 | 894513 | 894516 | 894512 | -0.1013647 |
| AGGTTCCTGGTATCAAAGTG | 7 | 767344 | 767363 | 767341 | 767344 | -0.0415362 |
| ATTCACATTCGAGGGGTTGC | 4 | 238808 | 238827 | 238830 | 238826 | -0.0738085 |
| AAAACACGTTTGCATTGTTT | 16 | 534884 | 534903 | 534906 | 534894 | -0.1368684 |
| TTAATCCGAGGTAATAGATT | 2 | 124958 | 124977 | 124955 | 124959 | -0.0926067 |
| GATAATATCACCGTAGTTTC | 13 | 99571 | 99590 | 99568 | 99579 | -0.1239932 |
| ATGCGTACAAGGAAGCGGGT | 2 | 186645 | 186664 | 186642 | 186648 | -0.0357824 |
| TAATAATTCTCAATAGATAC | 15 | 181592 | 181611 | 181614 | 181607 | -0.1913804 |
| GAACATCGATAAGTCGAATG | 13 | 275896 | 275915 | 275918 | 275910 | -0.09468 |
| TATTTCTCTGTTAGATGCTC | 15 | 633677 | 633696 | 633699 | 633695 | -0.1050648 |
| AGTTCCACTTTTCAGGCAGG | 5 | 196820 | 196839 | 196842 | 196832 | -0.0352892 |
| GTTACCCTAATCTATTACCT | 2 | 124950 | 124969 | 124972 | 124966 | -0.2212353 |
| TCGTAGGTTCGAATATCTCG | 4 | 1398891 | 1398910 | 1398913 | 1398909 | -0.0661702 |
| TTCACATTCGAGGGGTTGCC | 4 | 238809 | 238828 | 238831 | 238826 | -0.0751744 |
| CCAACTGCCTCAAAAAGGTA | 9 | 261058 | 261077 | 261055 | 261061 | -0.1216244 |
| ACAGCAAAAAATAAGACTGC | 15 | 802393 | 802412 | 802390 | 802400 | -0.1444372 |
| CCACTACATTTAGGTAGACG | 13 | 901618 | 901637 | 901640 | 901630 | -0.1003644 |

**Supplemental Table 3: Table of gRNAs that are likely to be affected by SNPs near their binding location.** The gRNAs shown all have significant background-dependent effects, a mapped QTL within 10 kb of the gRNA binding location, and a known BY/3S polymorphism at the gRNA binding location. More conservative thresholds were used when excluding gRNAs from analysis (see Methods section “Linkage Mapping”).

| gRNA Sequence | ORF | Chr | gRNA Position | Peak Start | Peak End | Peak Marker | -log10(pval) | condition |
|---|---|---|---|---|---|---|---|---|
| ATTAAAGTCATTTAGAGCAA | YBR237W | 2 | 691848 | 678574 | 705106 | 681557 | 18.46 | BOTH |
| AAAAGCACACCGTATAGAAA | YJR040W | 10 | 507680 | 497086 | 545773 | 526954 | 14.05 | BOTH |
| AAGGTTCCTGGTATCAAAGT | YGR138C | 7 | 767345 | 748014 | 779852 | 766068 | 13.75 | BOTH |
| TGAGTTCTAGCGGAAACCGC | YML098W | 13 | 77154 | 27643 | 73993 | 32621 | 6.38 | BOTH |
| TTTTGGCAAGCACGAATTGA | YJR111C | 10 | 636901 | 642106 | 677358 | 660402 | 5.2 | BOTH |
| GATACCAACCGATCCTTGCT | YGL167C | 7 | 190876 | 154206 | 226152 | 175590 | 5.22 | BOTH |
| TTCGCGTGGCATAACCCGGC | YML085C | 13 | 99549 | 57144 | 104510 | 93860 | 6.95 | BOTH |
| ATTTTGCTGGAAGGGCAATA | YLR347C | 12 | 826667 | 785227 | 846812 | 807311 | 6.66 | BOTH |
| AAGCTAATTAAGTGTAAACT | YKL028W | 11 | 385663 | 367086 | 420060 | 394572 | 5.35 | BOTH |
| ACTATCATAGGTATAAATAC | YKL137W | 11 | 185849 | 164020 | 193550 | 184014 | 8.21 | BOTH |
| ATGAGATTGAAACAATAACA | YJR112W | 10 | 636945 | 641813 | 683971 | 655475 | 5.94 | ATC2 |
| TACTGGCCGCCGGCATGCGA | YJR002W | 10 | 438744 | 415069 | 456781 | 434839 | 19.57 | BOTH |
| AGAAGTATGGATGGTTCGAA | YGR186W | 7 | 867710 | 846529 | 905027 | 869612 | 9.62 | BOTH |
| ACCAAGACGCCACTACATTT | YMR314W | 13 | 901609 | 899812 | 904006 | 900561 | 15.2 | BOTH |
| TTTTATGAAGACTGGAGAAC | YJR040W | 10 | 507664 | 496029 | 560037 | 545773 | 4.93 | BOTH |
| GGCGGAAGGAGAAAGTTTCA | YDL139C | 4 | 212162 | 176628 | 238826 | 222741 | 4.71 | ATC1 |
| TGTTAGTATTTAAGCTTATT | YMR287C | 13 | 845438 | 824397 | 894039 | 855635 | 5.56 | ATC1 |
| TAAAAGCACACCGTATAGAA | YJR040W | 10 | 507679 | 496029 | 544322 | 514236 | 10.77 | ATC1 |
| TATGAAGACTGGAGAACAGG | YJR040W | 10 | 507661 | 496029 | 571066 | 531791 | 6.36 | ATC1 |
| GTTAAGCATTGTAAAATACC | YPL173W | 16 | 222985 | 208747 | 244227 | 222598 | 4.5 | ATC1 |
| AAAGGTAAGAGATAATACTA | YOR159C | 15 | 633624 | 571939 | 652034 | 585188 | 5.96 | ATC1 |

**Supplemental Table 4: Table of gRNAs with a mapped QTL near their binding location and without an overlapping polymorphism.** The gRNAs shown all have background-dependent effects and a mapped QTL within 10 kb of the gRNA binding site. However, they lack any known polymorphism overlapping the gRNA binding site. All positions (gRNA position, interval start, interval end, and peak marker) are given in genome position. The width of the 2x-$\log_{10}$(pval) confidence interval is shown by the "Interval Start" and "Interval End" columns. The "Condition" column describes which replicates a mapped QTL was detected in.

| Hub Chromosome | Number of gRNAs | Hub Marker Position | Peak Width (bp) |
|---|---|---|---|
| VII | 38 | 979647 | 3371 |
| X | 25 | 646356 | 4918 |
| XIII | 76 | 43096 | 20469 |
| XIV | 18 | 470844 | 8643 |

**Supplemental Table 5: Positional information about hubs.** For each hub, the peak was defined as the marker of maximum significance across all gRNA interactions shown by a hub. The peak width was defined as the region where the highest number of confidence intervals across all gRNA interactions shown by a hub overlapped.

**Supplemental Data**

**Supplemental Data 1:** Barcode data for all segregants used in the fitness assay. Each row represents a unique segregant. The columns “seg_name”, “geno_bc”, and “genotype” describe the segregant name, segregant barcode, and segregant genotype, respectively.

**Supplemental Data 2:** Genotype data for all segregants used in the fitness assay. Rows “c”, “p”, and “gp” correspond to chromosome, position, and genome position. Each row is a polymorphic site among the segregants used in the assay. Each column contains the genotype data for a single segregant, with 0 indicating a BY allele and 1 indicating a 3S allele at that locus.

**Supplemental Data 3:** Data for all gRNAs used in the fitness assay. The “guide”, “category”, “chrom”, “strand”,  “start_gp”, and “end_gp” columns describe the guide sequence, guide library of origin, target chromosome, target strand, 5’ genome position, and 3’ genome position, respectively. The “pam.mid.gp”, “ORF”, and  “Gene_name” describe the midpoint of the PAM, the systematic name of the targeted ORF, and the standard name of the targeted ORF (if applicable). Control gRNAs show “NA” under the ORF column.

**Supplemental Data 4:** Barcodes recovered per segregant. Each row is a unique segregant. The “genotype” column provides the segregant name. The “atc1_barcodes”, “atc2_barcodes”, and “con3_barcodes” describe the number of unique double barcodes recovered via sequencing for that segregant. The “atc1_normalized_reads”, “atc2_normalized_reads”, and “con3_normalized_reads” describe the number of normalized reads recovered for that segregant in each fitness assay.

**Supplemental Data 5:** Barcodes recovered per gRNA. Each row is a unique gRNA, with the “guide” column providing the gRNA name. The “atc1_barcodes”, “atc2_barcodes”, and “con3_barcodes” describe the number of unique double barcodes recovered via sequencing for that gRNA. The “atc1_normalized_reads”, “atc2_normalized_reads”, and “con3_normalized_reads” describe the number of normalized reads recovered for that gRNA in each fitness assay. The “number_of_genotypes” column shows how many segregants that gRNA appeared in.

**Supplemental Data 6:** Processed data from the ATC1 fitness assay. Each row represents a unique double barcode lineage. All double barcodes are linked to their respective segregant or gRNA. Each column contains the following information:
geno_bc: The genotype barcode sequence.
guide_bc: The gRNA barcode sequence.
T0: Normalized read count at time point 0 for this assay. Multiple adjacent columns follow this format (i.e. T1 for time point 1).
fitness: Fitness estimate from PyFitSeq software.
error: Reported error for the fitness estimate from PyFitSeq software.
Log_likelihood: Log likelihood score of the fitness estimate from PyFitSeq software.
Seg_name: Segregant name associated with the corresponding segregant barcode.
Genotype: Genotype name associated with that segregant.
Guide_seq: gRNA sequence associated with the gRNA barcode.
Gene_orf: The systematic name of the ORF targeted by the gRNA. This is “NA” for control guides.
Gene_name: The standard name of the ORF targeted by the gRNA, if applicable.
Guide_category: The plasmid library the gRNA originates from, with “E” indicating the essential set used for analysis.

**Supplemental Data 7:** Processed data from the ATC2 fitness assay. Formatting is identical to file Supplemental Data 6.

**Supplemental Data 8:** Processed data from the CON fitness assay. Formatting is identical to file Supplemental Data 6.

**Supplemental Data 9:** Effects of gRNAs. Each row is a unique gRNA, with the "guide", "gene_orf", and "gene_name" providing the gRNA sequence, systematic name of the targeted ORF, and standard name of the targeted ORF. The "main.p" column is the p-value of the *gRNA* term from the linear model , and "int.p" is the p-value of the *genotype:gRNA* interaction term. The "main.fdr" and "int.fdr" columns are the same p-values after Benjamini-Hochberg multiple testing correction. The "nd.main.effect" columns shows if that gRNA had a mean effect three standard deviations below the median of the simulated distribution of neutral guides. The "guide.model.coef" column is the coefficient of the *gRNA* term extracted from the appropriate linear model. The "number.of.points" and "number.of.genos" columns show how many double barcodes and segregants were used to calculate guide effects, respectively.

**Supplemental Data 10:** Deviation values of gRNAs. Each column in this table is a unique gRNA, and each row is a unique segregant. The first column is composed of segregant names. Only gRNAs with background effects are present.

**Supplemental Data 11:** Results of linkage mapping on deviation values. Each row is a unique 2x-$\log_{10}$(pval) drop confidence interval. The "chr", "guide", and "ORF" columns indicate the chromosome target, gRNA sequence, and systematic name of the targeted ORF. The "peak.start" and "peak.end" columns indicate the genome positions of the 5' end and 3' end of the confidence interval. The "peak.marker", "lod.score", "adj_r2", and "locus_coef" show the genome position of the peak marker, 2x-$\log_{10}$(pval), variance explained, and coefficient of the *locus* term, all from the linear model used at the peak marker locus. The columns "total.geno.used" and "locus_id" describe the number of unique segregants used when running the linear model, as well as the hub the interval is associated with (if any).

**Supplemental Data 12:** Genotype data for all segregants in the entire haploid panel. Formatting is identical to Supplemental Data 2.

**Supplemental Data 13:** Table of all SNPs present in the 3S strain, relative to S288C. Column 1 is the chromosome, column 2 is the SNP position, column 3 is the nucleotide in S288C, and column 4 is the nucleotide in 3S. The S288C reference differs from the BY strain by only roughly 125 SNPs.

**Supplemental Data 14:** Table of all linkages between gRNA barcodes and gRNA sequences. Column 1 is the gRNA barcode, column 2 is the gRNA sequence, column 3 is the targeted ORF, column 5 is the common name of the targeted gene, and column 5 is the classification of the targeted gene (essential or non-essential).
